# Protective effects of saffron and its constituent crocetin to motor symptoms, short life span and rough-eyed phenotypes in the fly models of Parkinson’s disease *in vivo*

**DOI:** 10.1101/2020.12.06.408997

**Authors:** Eiji Inoue, Takahiro Suzuki, Yasuharu Shimizu, Keiichi Sudo, Haruhisa Kawasaki, Norio Ishida

## Abstract

Parkinson’s disease (PD) is a common neurodegenerative disorder with motor symptoms linked to the loss of dopaminergic neurons in the brain. α-Synuclein is an aggregation-prone neural protein that plays a role in the pathogenesis of PD. In our previous paper, we found that saffron; the stigma of *Crocus sativus* Linné (*Iridaceae*), and its constituents (crocin and crocetin) suppressed aggregation of α-synuclein and promoted the dissociation of α-synuclein fibrils *in vitro*. In this study, we investigated the effect of dietary saffron and its constituent, crocetin, *in vivo* on a fly PD model overexpressing several mutant α-synuclein in a tissue-specific manner. Saffron and crocetin significantly suppressed the decrease of climbing ability in the *Drosophila* overexpressing A30P (A30P fly PD model) or G51D (G51D fly PD model) mutated α-synuclein in neurons. Saffron and crocetin extended the life span in the G51D fly PD model. Saffron suppressed the rough-eyed phenotype and the dispersion of the size histogram of the ocular long axis in A30P fly PD model in eye. Saffron had a cytoprotective effect on a human neuronal cell line with α-synuclein fibrils. These data showed that saffron and its constituent crocetin have protective effects on the progression of PD disease in animals *in vivo* and suggest that saffron and crocetin can be used to treat PD.

## Introduction

Parkinson’s disease (PD) is a common neurodegenerative disorder characterized by abnormal accumulation of α-synuclein aggregates (called Lewy bodies) in the dopaminergic neurons of the substantia nigra and other brain regions (1–3). The aggregation of α-synuclein promotes dopaminergic neuron dysfunction and death, which is associated with the pathogenesis and progression of PD (4–6). PD causes motor symptoms such as resting tremor, rigidity, bradykinesia, and postural instability (7). Replacement therapy with levodopa, a dopamine precursor, is effective against several symptoms of PD (8). However, when PD progresses and more neurons die, levodopa becomes less effective. Furthermore, levodopa causes a complication characterized as involuntary writhing movements (9, 10). Therefore, new therapeutic agents for PD are required.

Saffron is the stigma of *Crocus sativus* Linné (*Iridaceae*). It is both a food additive (used as a spice, yellow food coloring, and a flavoring agent) and a drug (used as an analgesic, sedative, and emmenagogue) (11). Saffron contains two main carotenoids, crocetin and crocin (crocetin digentiobiose ester) (12), and they inhibit amyloid-β aggregation *in vitro* (13–15). Our previous paper showed that saffron and its constituents (crocetin and crocin) inhibit α-synuclein aggregation and dissociate the formed aggregate *in vitro* (16). Ghasemi *et al.* reported that crocin inhibited α-synuclein aggregation *in vitro* and explored its mechanism *in silico* (17). These papers indicate that saffron may prevent the accumulation of aggregated α-synuclein in PD neuronal cells *in vivo*. Many animal PD models exist (18). Among them, many researchers have opted for transgenic *Drosophila* models, as these non-mammalian animal models are suitable for fast, high-throughput drug screening (19–22). In this study, we investigated the effect of saffron and its constituent crocetin on the *Drosophila PD model in vivo.* The data indicate that saffron and crocetin might be useful to treat PD, because they ameliorate the phenotype of abnormal α-synuclein in eyes and motor symptoms, short life span of PD model flies.

## Results

### Saffron and crocetin suppressed the decrease of climbing ability in *Drosophila* expressing A30P or G51D mutated α-synuclein

A30P or G51D flies have a lower climbing ability than normal flies (23, 24). Saffron and crocetin significantly suppressed the decrease of climbing ability in the A30P flies (Fig. 1A, 1B) or G51D flies (Fig. 1C, 1D).

**Fig. 1.**
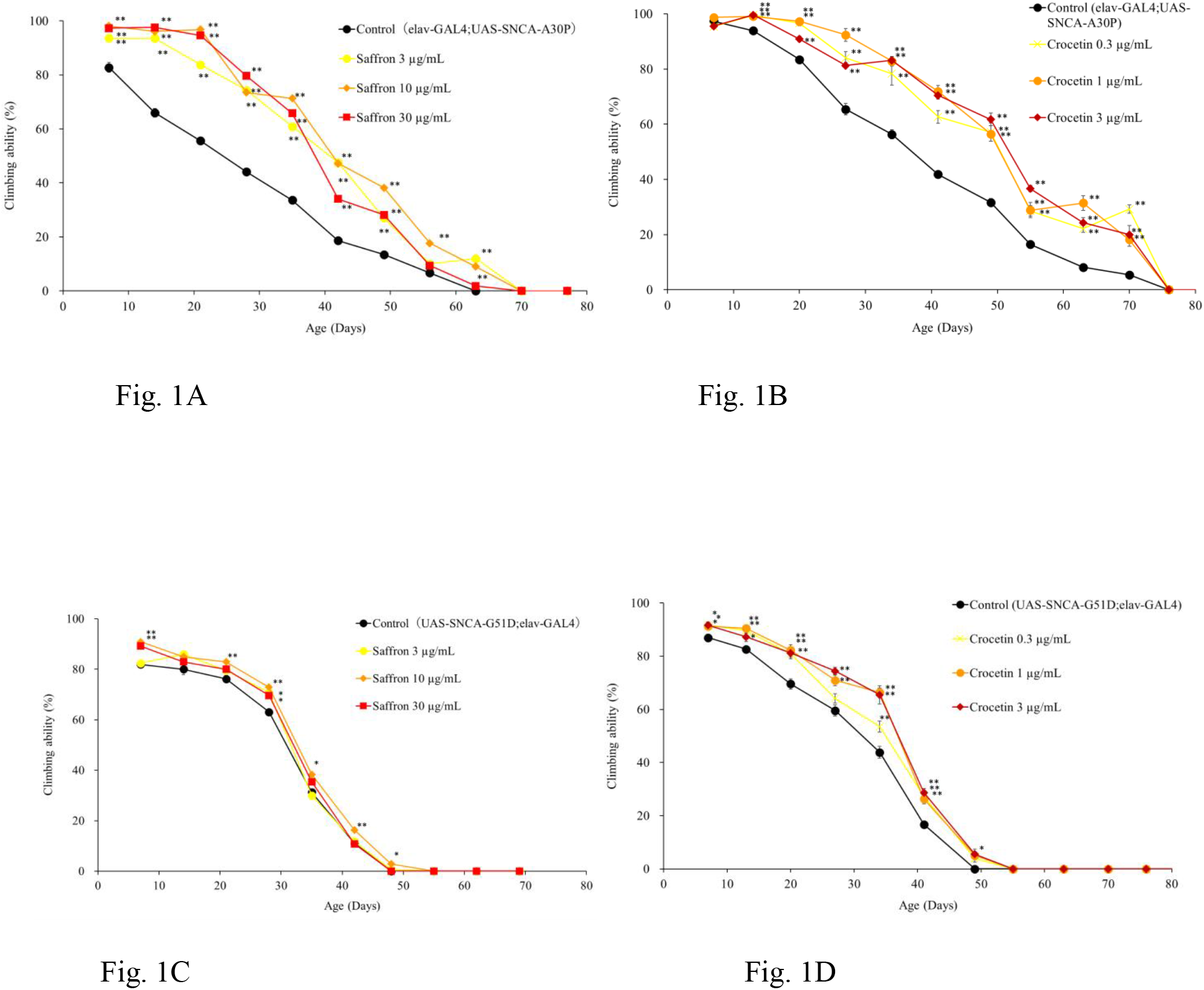
Saffron and crocetin suppressed the decrease of climbing ability in A30P or G51D flies. (A, B) Saffron and crocetin suppressed the decrease of climbing ability in A30P flies. (C, D) Saffron and crocetin suppressed the decrease of climbing ability in G51D flies. Data are expressed as the mean ± SEM of ten trials at each point. (N = 24–30). **: P < 0.01, *: P < 0.05 as compared with the control group using Dunnett’s test.

### Saffron and crocetin extended the life span of the G51D flies

There is little life span difference between A30P flies and normal flies (23), whereas G51D flies have a shorter life span than normal flies as expected (24). Saffron and crocetin did not affect the survival rate of the A30P flies (Fig. 2A, 2B). In contrast, three different concentrations of saffron and 3 μg/ml crocetin significantly increased the survival rate of the G51D flies (Fig. 2C, 2D).

**Fig. 2.**
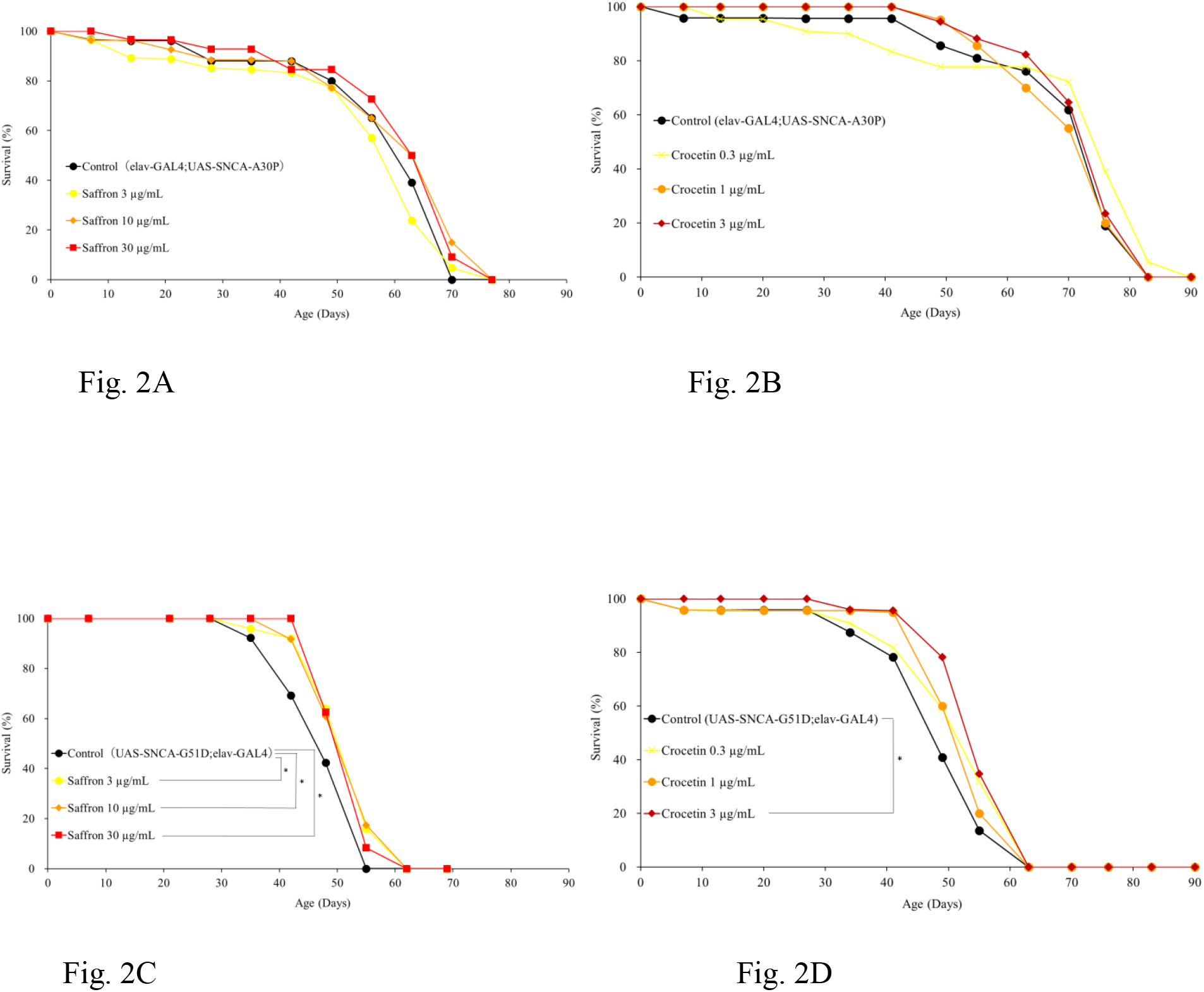
Saffron and crocetin extended the life span of the G51D flies. (A, B) Saffron and crocetin did not affect the life span of the A30P flies. (C, D) Saffron and crocetin extended the life span of the G51D flies. The curves show the survival rates of the flies. (N = 24–30). *: P < 0.05 as compared with the control group using the log-rank test.

### Saffron suppressed the eye morphological defects in the A30P flies

The A30P flies showed a rough-eyed phenotype (Fig. 3A). The size histogram of the major axis of the ommatidium of the A30P flies was more dispersed than that of the normal flies. Saffron significantly suppressed the eye morphological defects in the A30P flies, and the effect was remarkable at 30 μg/mL (Fig. 3B).

**Fig. 3.**
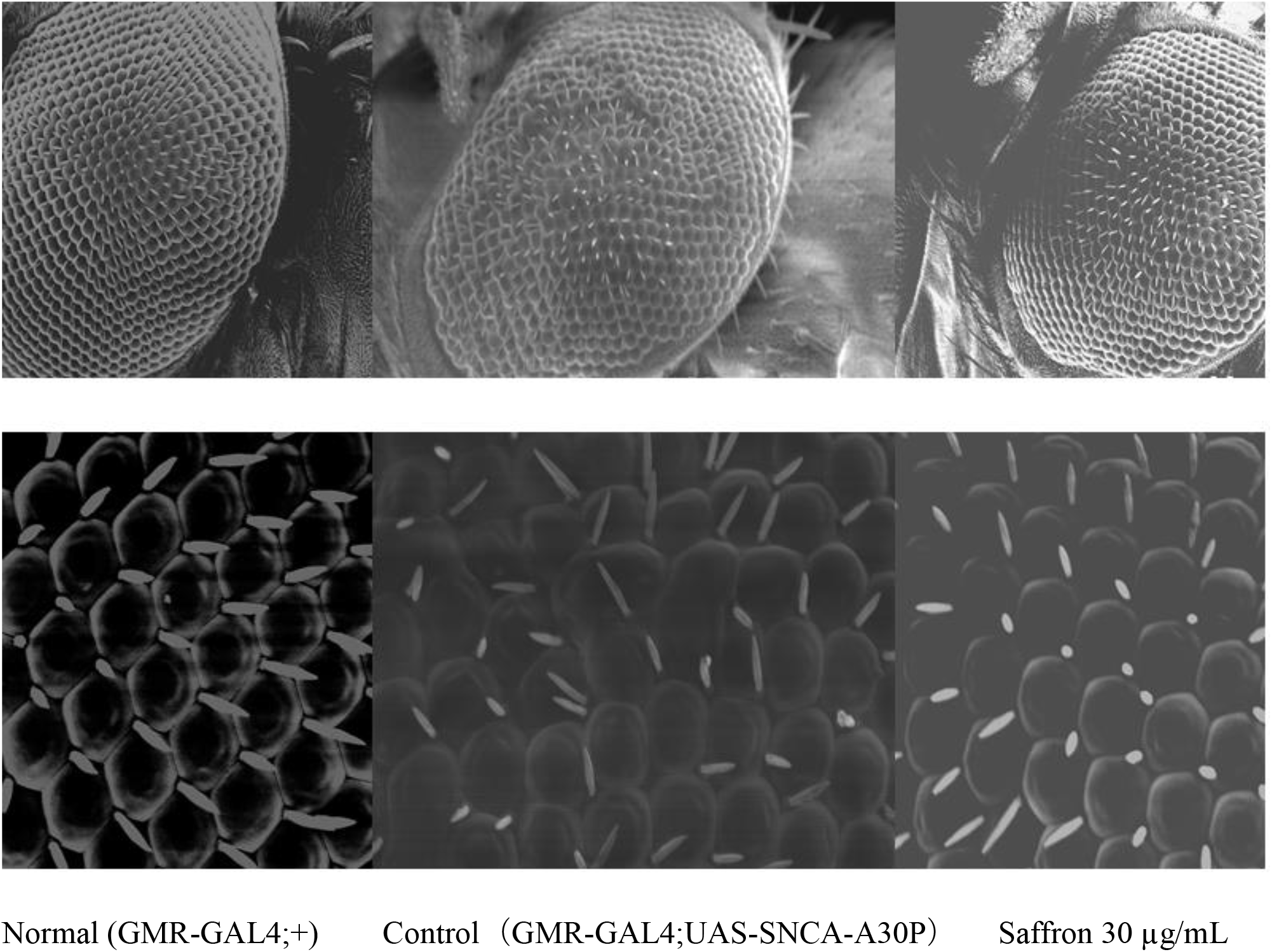
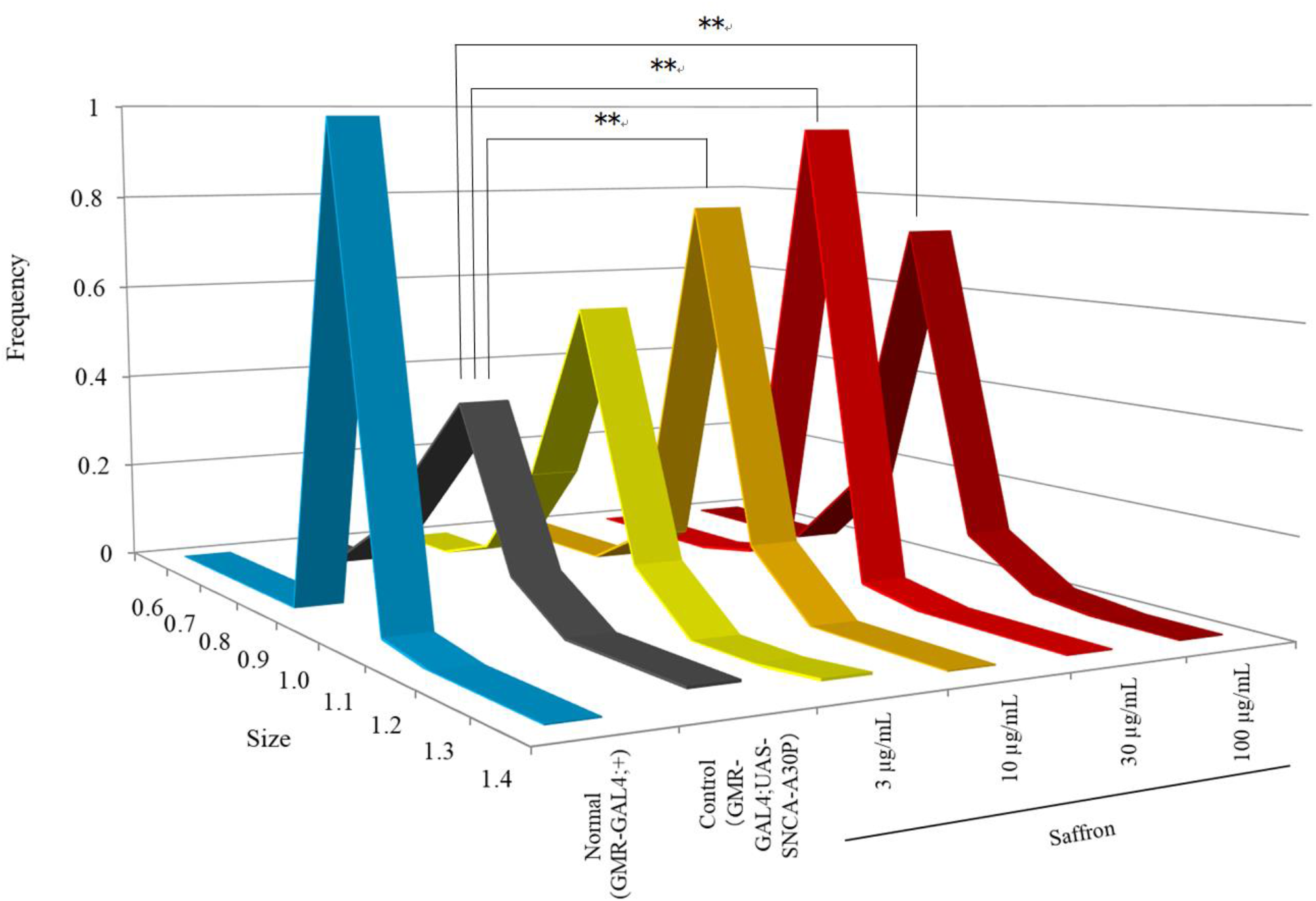
Saffron suppressed the eye morphological defects in the A30P flies. (A) The A30P flies had a rough-eyed phenotype and saffron suppressed the phenotype expression. Scale bars indicate a length of 100 (Upper) or 10 (lower) μm. (B) Size histogram of the major axis of the ommatidium in the A30P flies compared with that of the normal flies. **: P < 0.01 as compared with the control group using Levene’s test. (N = 6 on Normal, N = 7 on Control, N = 8 on saffron 3 μg/mL, N = 3 on saffron 10 μg/mL, N = 9 on saffron 30 μg/mL, N = 8 on saffron 100 μg/mL).

### Saffron had a cytoprotective effect on SH-SY5Y cells cultured with α-synuclein fibrils

PD research is mostly conducted on human neuronal cell models, such as the neuroblastoma SH-SY5Y cells. The α-synuclein fibrils-treatment decre**ased** SH-SY5Y cells viability by 23%. Saffron had a cytoprotective effect, and the effect was remarkable at 20 μg/mL (Fig. 4).

**Fig. 4.**
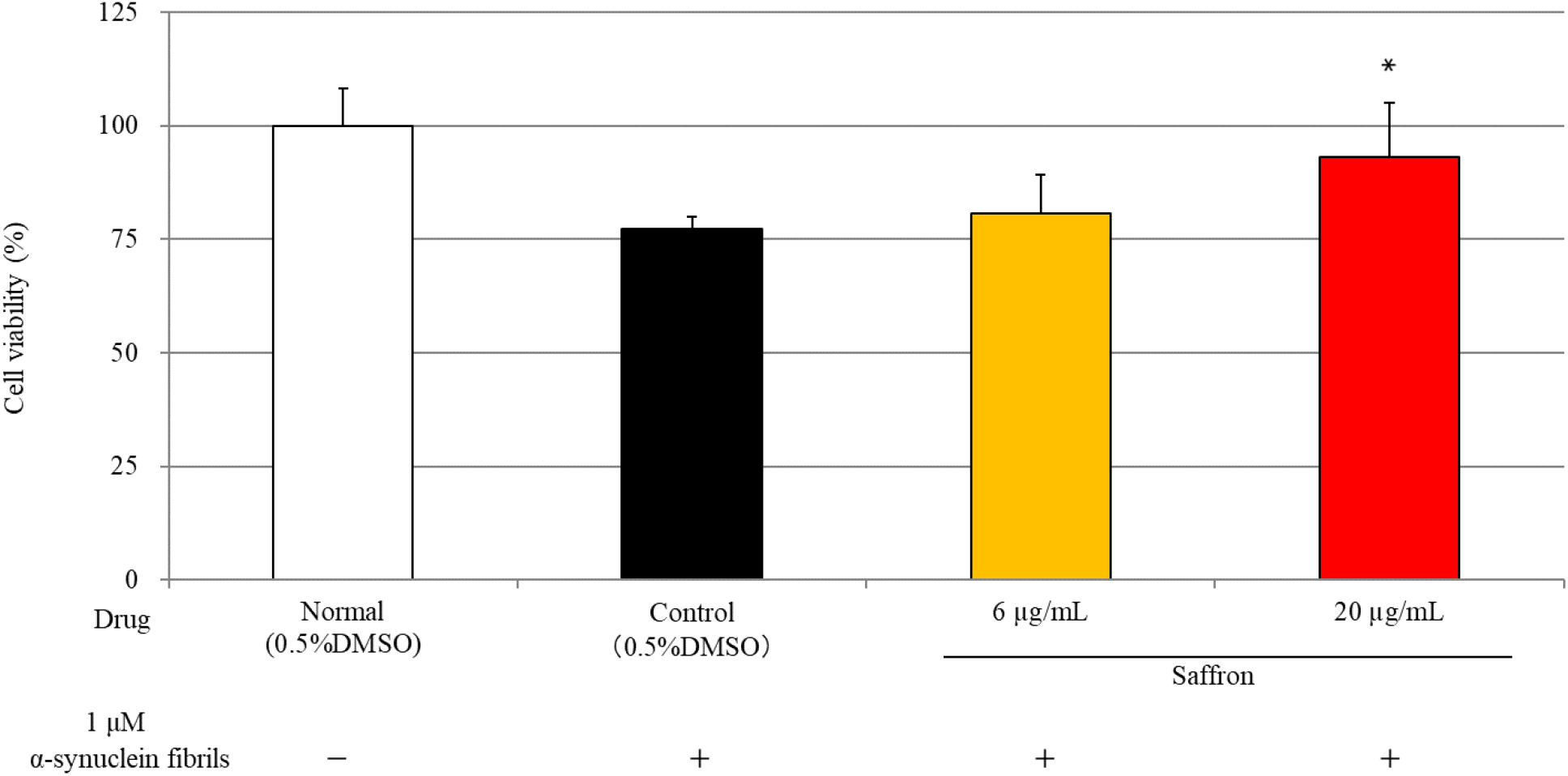
Saffron had a cytoprotective effect on SH-SY5Y cells cultured with α-synuclein fibrils. Data are expressed as the mean ± SEM. (N = 6 or 3 on saffron 6 μg/mL). *: P < 0.05 as compared with the control group using Dunnett’s test.

## Discussion

To investigate the effect of saffron on *in vivo* animal PD models, we used transgenic *Drosophila* models via the UAS/GAL4 system. We transfected A30P or G51D predominantly in the neurons of the flies using the elav driver (elav-GAL4;UAS-SNCA-A30P or UAS-SNCA-G51D;elav-GAL4 strain) and A30P in the eyes of the flies using the GMR driver (GMR-GAL4;UAS-SNCA-A30P strain). These models, which overexpress α-synuclein in the neurons or eyes show an age-dependent decrease in climbing ability, shorter life span, or retinal degeneration (19–21, 23–25) and are used to screen for drugs effective on PD (19–21). Besides, the A30P and G51D fly PD models have more severe PD-like symptoms than wild-type fly PD models (23–24). Our previous paper indicated that an *in vitro* saffron treatment inhibited the formation of α-synuclein aggregates in a dose-dependent manner (16). The data suggest that saffron ameliorates these abnormal phenotypes in the PD fly models. Saffron has multiple pharmacological properties, such as a potent antioxidant activity (26). The accumulation of Lewy bodies due to overexpression of α-synuclein generates reactive oxygen species (ROS) (27). ROS create a state of oxidative stress and damage the neurons including the dopaminergic neurons (27). Besides, ROS generation causes locomotor dysfunction in flies (20). These papers suggest that saffron’s antioxidative effect is involved in the effects on PD fly models.

Herein, we found that crocetin had a similar protective effect to saffron on PD fly models. Among the constituents of saffron, we have reported that crocin and crocetin prevent the formation of α-synuclein aggregates and dissociate the formed aggregates *in vitro* (16). Although saffron contains an overwhelmingly higher amount of crocin than crocetin (16), this compound hardly crosses the blood-brain barrier. However, crocetin crosses the blood-brain barrier by oral administration (28), and is formed from crocin hydrolysis in the gastrointestinal tract and epithelium in animals (29). These facts suggest that the effects of saffron on the PD fly models are partially caused by crocetin which is converted from crocin in the animal body.

Saffron had a cytoprotective effect on SH-SY5Y cells cultured with α-synuclein fibrils. Current PD research is mostly conducted on human neuronal cell models, such as the neuroblastoma SH-SY5Y lineage (30), because human dopaminergic neurons, the cells mainly affected in PD, are difficult to obtain and maintain as primary cells. This cell line is chosen because of its human origin and catecholaminergic neuronal cell properties (30). Our results reveal that saffron can prevent the cytotoxicity of α-synuclein fibrils on human-derived cells.

In conclusion, we showed that saffron modulated the PD-like phenotypes of *Drosophila* overexpressing α-synuclein *in vivo*, and its constituent crocetin had similar effects. This suggests that saffron and crocetin modulate the phenotypes by inhibiting the α-synuclein aggregate formation and dissociating the formed aggregates (Fig. 5). Further studies on the molecular mechanisms are required to clarify the effects of saffron and crocetin on PD.

**Fig. 5.**
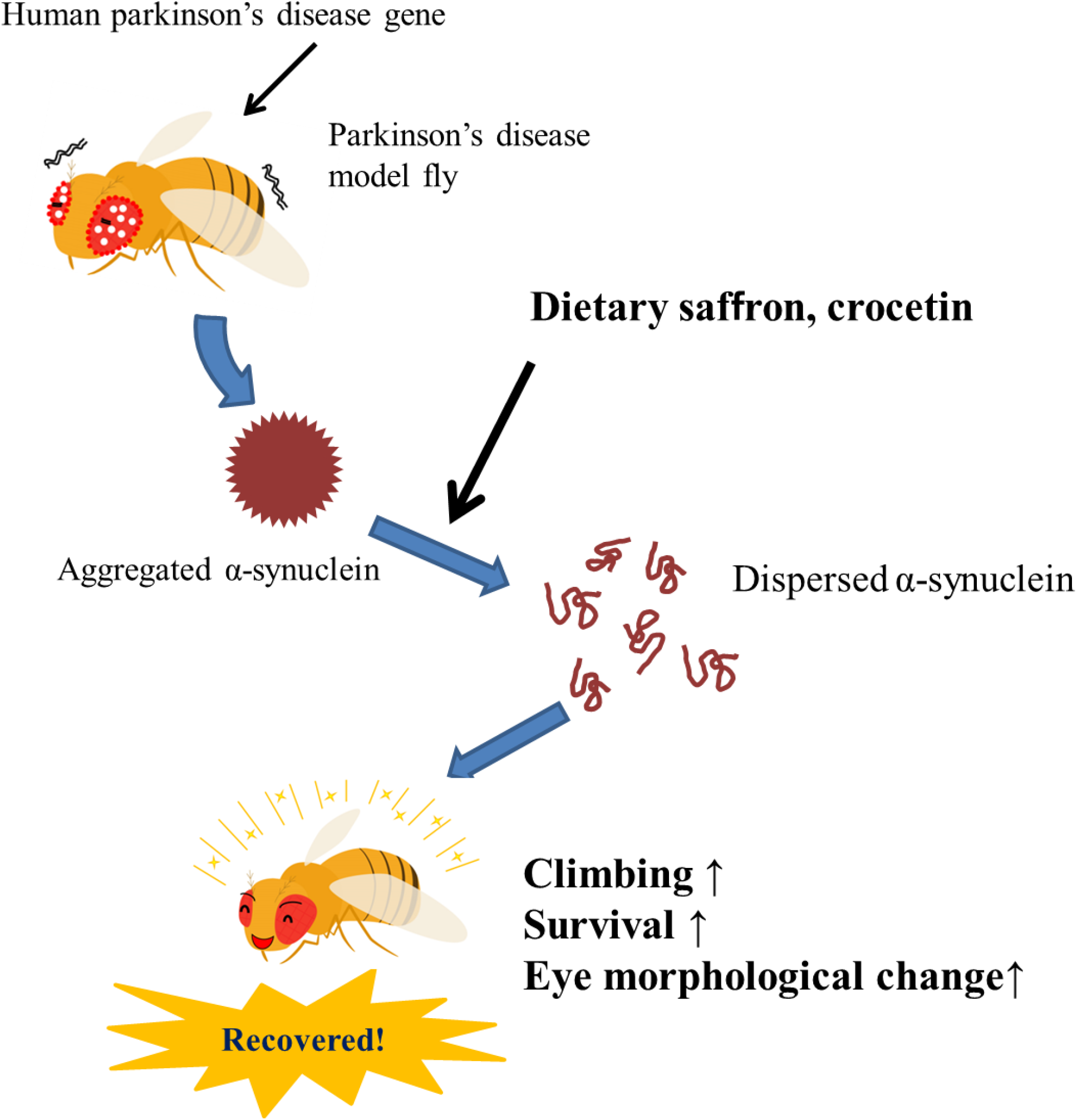
Graphical concepts. Human Parkinson’s disease genes (A30P or G51D mutated α-synuclein) were expressed in the flies’ eyes or neurons using the elav or GMR drivers, respectively. Administering dietary saffron and crocetin to PD model flies had a positive effect on α-synuclein aggregation, climbing ability, survival rate, and eye morphological defects.

## Materials and Methods

### Reagents

Saffron was obtained from Matsuura Yakugyo (Aichi, Japan). For *in vivo* studies, saffron was suspended in distilled water and added to the *Drosophila* food. For *in vitro* studies, the saffron extract was prepared with 10% dimethyl sulfoxide (DMSO)/Dulbecco’s phosphate-buffered saline (D-PBS) (9.5 mM, pH 7.4) using an ultrasonic generator (Model 2510, Branson, CT, USA) for 30 min at 25°C. The extract was centrifuged (3000 rpm, 10 min), the supernatant was sterilized by filtration using a 0.2 μm pore size membrane filter (25CS020AS, Advantec Toyo, Tokyo, Japan) and added to the culture medium. Acetonitrile, Dulbecco’s Modified Eagle Medium (DMEM)/F-12 medium, D-PBS, and DMSO were purchased from FUJIFILM Wako Pure Chemical Industries (Osaka, Japan). The α-Synuclein fibrils were purchased from Cosmo Bio (Tokyo, Japan). The MEM non-essential amino acids solution was purchased from Nacalai Tesque (Kyoto, Japan). The penicillin-streptomycin solution was purchased from Sigma-Aldrich (St. Louis, MO, USA). Fetal bovine serum (FBS) was purchased from Life Technologies (Grand Island, NY, USA). Cell Titer 96^®^ was purchased from Promega (Madison, WI, USA).

### *Drosophila* stocks, crosses and maintenance

The UAS-SNCA-A30P, UAS-SNCA-G51D, and elav-GAL4/CyO strains were obtained from the Bloomington Drosophila Stock Center (Bloomington, IN, USA). The IF/CyO;elav-GAL4 strain was kindly provided by Dr. Masami Shimoda from The National Agriculture and Food Research Organization (NARO). The GMR-GAL4 strain was obtained from the National Institute of Genetics Fly Stock Center (Shizuoka, Japan).

Transgenic females carrying UAS-SNCA-A30P were crossed with males carrying elav-GAL4/CyO to generate the *Drosophila* PD model expressing human A30P mutated α-synuclein in neurons. Transgenic females carrying UAS-SNCA-G51D were crossed with males carrying IF/CyO;elav-GAL4 to generate the *Drosophila* PD model expressing human G51D mutated α-synuclein in neurons. Transgenic females carrying UAS-SNCA-A30P were crossed with males carrying elav-GAL4/CyO to generate the *Drosophila* PD model expressing human A30P mutated α-synuclein in the eyes.

The *Drosophila* strains were maintained as previously described (31). *Drosophila* were reared in vials of standard yeast cornmeal at 25°C under a 12 h light/dark cycle at constant humidity (60%–80%). The *Drosophila* medium was prepared as previously described (31). The boiled standard medium consisting of 8% cornmeal, 5% glucose, 5% dry yeast extract, and 0.64% agar was supplemented with 0.5% propionic acid and 0.5% butyl p-hydroxybenzoate. The crossings, survival assays, and climbing ability assays were conducted at 25°C. The eye analysis experiment was conducted at 29°C.

### Climbing assay and life span

Male flies of the elav-GAL4;UAS-SNCA-A30P and UAS-SNCA-G51D;elav-GAL4 strains were collected upon eclosion and divided into multiple groups of 24–30 individuals each. The flies were housed at a density of 8–10 flies in a vial and transferred to another vial with fresh medium every three or four days.

The climbing assay was conducted as described previously (23–24). Briefly, the flies were placed in an empty polystyrene vial (9.5 cm × 2.5 cm). The flies were placed at the bottom of the vial by gentle tapping, and the flies that climbed and crossed the 8-cm high mark in the 10 s following the tapping were counted. The test was repeated 10 times for each set of flies.

The *Drosophila* were observed during this experiment, and the survival rate of each group was calculated once a week. The flies were considered dead when they did not move despite agitation (25, 32).

### *Drosophila* eye imaging

The UAS-SNCA-A30P and GMR-GAL4 strains were crossed at 29°C in the drug-containing medium. Then the three-day-old male flies of the GMR-GAL4;UAS-SNCA-A30P strain were collected and their head was mounted on a stage with double-sided tape and sputter-coated with gold. Scanning electron microscopic images were taken with a JSM-6301F electron microscope (JEOL, Tokyo, Japan) at 5 kV using a previously described protocol (33–34).

### Cytotoxicity Assay

SH-SY5Y cells were purchased from DS Pharma Biomedical (Osaka, Japan) and maintained in DMEM/F-12 medium with 10% FBS, 1% MEM non-essential amino acids solution, 100 U/mL penicillin, and 100 μg/mL streptomycin at 37°C in 5% CO_2_. The cells were cultured in 96-well microplates (353072, Corning, NY, USA) at a density of 1 × 10^4^ cells/well for 24 h, then the medium was replaced with a new medium containing each drug and 1 μM of α-synuclein fibrils and the cells were incubated for 48 h. Cytotoxicity was calculated by an MTS assay using CellTiter 96^®^. The absorbance of the formazan dye in the well at 490 nm was measured using a microplate reader (Synergy H1, Biotec, VT, USA).

### Statistical analysis

Data were analyzed using Excel (Microsoft, Redmond, WA, USA), KaleidaGraph (HULINKS, Tokyo, Japan), or R (The R Foundation for Statistical Computing, Vienna, Austria). Data from the climbing assay and cytotoxicity assay were analyzed using One-way ANOVA followed by Dunnett’s post-hoc test. Data from the survival rate was analyzed using the Logrank test. The variance of the major axis sizes of the ommatidium was analyzed using Dispersion analysis (Levene’s test).

## ACKNOWLEDGMENTS

This research was supported by Kyushin Pharmaceutical Co., Ltd. We also thank Dr. Masami Shimoda from NARO for help with electron microscopy experiments and a generous gift of the *Drosophila* stocks.

## Footnotes

The authors declare no competing interest.

